# Cross-Species Translation Enhances the Use of Mouse Models for Translatability and Drug Discovery in Late-Onset Alzheimer’s Disease

**DOI:** 10.64898/2026.03.21.713391

**Authors:** Jee Hyun Park, Jinsheng Yu, Brendan P. Lucey, Douglas K. Brubaker

## Abstract

Alzheimer’s disease (AD) is a brain disease characterized by deposition of insoluble amyloid-β plaque, intraneuronal neurofibrillary tangles, and cognitive dysfunction. AD can be characterized as early-onset or late-onset based on age and genetic factors. For early-onset, these genetic factors can include amyloid precursor protein (APP), presenilin-1 (PSEN1), and presenilin-2 (PSEN2). For late-onset, these can include apolipoprotein E e4 (APOE4), and the R47H variant of triggering receptor expressed on myeloid cells 2 (TREM2). Mouse models incorporating these risk factors provide critical knowledge for studying AD pathology and preclinical studies for drug development. However, these transgenic mice depend on early-onset genetic mutations and are deficient in certain AD features that are present in late-onset. Here, we developed innovative non-linear and feature selection procedures for our cross-species translation framework, Translatable Components Regression (TransComp-R), to identify transcriptomic features in mouse models predictive of human late-onset AD pathobiology. We used the cross-species computational translatability links of TransComp-R to perform computational high-throughput drug screening and identified multiple repurposable drugs for AD treatment that targeted the sleep-wake cycle. We tested these predictions in an orthogonal, prospective cohort of human subjects treated with an orexin receptor antagonist, suvorexant. We correlated conserved protein-level biomarkers from our cross-species transcriptomics model with significant reductions in phosphorylated tau in cerebrospinal fluid collected from humans treated with suvorexant. This study demonstrates the power of computational methods like TransComp-R to enhance the utility of murine disease models for discovering new therapeutic approaches for AD.

**One Sentence Summary:** Cross-species translation modeling across different mouse models reveals sleep-relevant drug mechanisms as potentially therapeutic for Alzheimer’s disease.

## INTRODUCTION

Alzheimer’s disease (AD) is a neurodegenerative disease with over 6 million cases in the United States (*1*). People with AD often display compromised memory, alterations in mood, and behavioral changes throughout disease progression (*2*). In addition, AD involves the deposition of insoluble amyloid-β (Aβ) plaques, intraneuronal neurofibrillary tangles of tau, synaptic and neuronal loss, cognitive dysfunction, as well as active microglia and astrocytes (*3–6*). Early-onset AD cases involve mutations in presenilin-1 (PSEN1), and presenilin-2 (PSEN2), and amyloid precursor protein (APP) (*7*). Age is the greatest risk factor for AD and most AD is late-onset in individuals 65 years and old (*8*). Apolipoprotein E e4 (APOE4) is the greatest genetic risk factor, increasing risk by 3x-12x for single or double APOE4 carriers, respectively (*9*). Another risk factor is a heterozygous mutation in triggering receptor expressed on myeloid cells 2 (TREM2) (*10*). The TREM2 R47H variant impairs the function of microglia and poses a risk equivalent to APOE4 (*11*, *12*). Furthermore, microglia and astrocytes, the immune cells in the central nervous system, induce neuroinflammation, which contributes to AD production of inflammatory cytokines, chemokines, and reactive oxygen species detrimental to synapses and neurons (*13*).

Multiple mouse models have been developed to study different aspects of AD pathology and preclinical intervention testing. Many mouse models are transgenic and express mutations linked to early-onset AD (*14*). These models lack major AD hallmarks, such as APP transgenic mice that develop amyloid plaques but not neurofibrillary tangles, and have an accelerated disease progression (*15*). Knock-in mouse models provide additional valuable insights into other pathways, especially for genetic factors associated with late-onset AD (*14*). Nevertheless, therapies tested in these mouse models frequently lack efficacy in patients and this lack of translatability remains a burden that limits the utility of animal models for the study of AD (*16*).

Our group has developed computational approaches designed to improve the predictability of therapeutic translation across species that have previously produced experimentally testable and validated predictions of therapeutic efficacy in humans that may be highly relevant to addressing the translatability challenge of mouse models in AD (*17*, *18*).

Here, we developed a non-linear feature selection-based implementation of our cross-species translation framework, Translatable Components Regression (TransComp-R), to determine the AD mouse model transcriptomic signatures predictive of AD pathology in humans. We used the computational mapping from mouse-to-human biology to conduct an *in silico* drug screening analysis to identify translatable therapeutic avenues for AD. Finally, we evaluated a candidate therapy modulating sleep-based mechanisms and phosphorylated tau levels in an independent prospective human cohort and measured protein-level biomarkers identified in our cross-species model. Overall, our study establishes the use of cross-species modeling to investigate translatable molecular characteristics and identify translatable therapeutic modalities for Alzheimer’s disease.

## RESULTS

### Generating robustly predictive cross-species transcriptomic models of AD phenotypes

We obtained murine gene expression data from GSE141509 on Gene Expression Omnibus (GEO) with samples from the brain hemispheres, including APOE4 KI (n=21), APOE4Trem2 (n=24), Trem2*R47H (n=23), 5xFAD (n=11), and control C57BL/6J (n=23) for 6- and 12-month-old mice (*19*). The mouse platform consisted of 760 NanoString probes determined by human co-expression modules of enriched and orthologous genes that are relevant in AD (*19*). For humans, we used GSE44772, and filtered for postmortem prefrontal cortex samples from late-onset AD (n=128) and non-dementia (n=58) subjects with ages greater than 60 years (*20*). We converted the mouse genes into human homologous genes and found overlapping genes between the mouse and human for downstream analysis (604 genes).

For our cross-species TransComp-R model, we performed principal component analysis (PCA) on each of the mouse model datasets for each age category and extracted the PCs explaining ∼80% cumulative variance (Table 1). Next, we conducted matrix multiplication to project the human transcriptomic samples onto the mouse transcriptomic PCs, thereby contextualizing human AD-associated variation in terms of murine model biology by relative separation in PC space **(Fig. 1).**

**Fig. 1.**
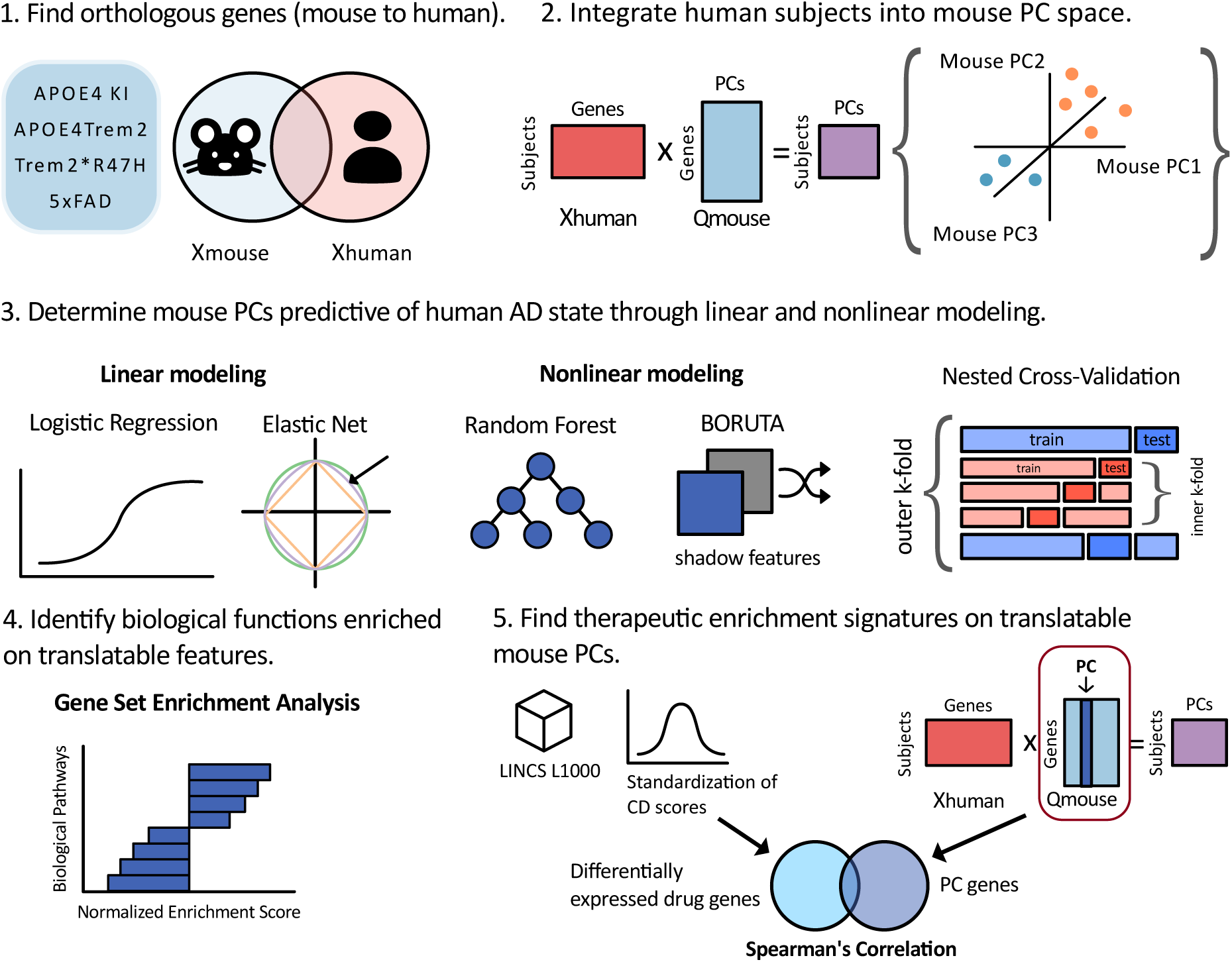
Pipeline for mouse to human computational translation. Methods used for our analysis, including building the TransComp-R model, modeling using linear and nonlinear methods, gene set enrichment analysis of the selected mouse features, and *in silico* drug screening analysis.

**Table 1.** Number of PCs for each mouse model. The number of principal components selected, explaining ∼80% variance for each mouse model type with respective control group by age.

| Age (months) | APOE4 KI (n) | APOE4Trem2 (n) | Trem2*R47H (n) | 5xFAD (n) |
| --- | --- | --- | --- | --- |
| 6 | 13 | 12 | 13 | 6 |
| 12 | 12 | 14 | 13 | 6 |
| 6 and 12 | 20 | 22 | 22 | 10 |

We implemented linear (logistic regression, elastic net) and nonlinear modeling (random forest, Boruta feature selection) methods to build models using human projections on murine PCs to predict human AD status and identify translatable biology **(Fig. 1).** All computational models performed comparatively well with high averaged area under the curves (AUC) of the 10 outer folds across the different mouse types (AUC range= 0.8906-0.9405) **(Fig. 2).** We found that mouse models combining 6- and 12-months data outperformed those using either age group alone; we therefore used the combined models for downstream analysis **(Fig. S1).** For mouse models, 5xFAD (avg. AUC = 0.930) had the highest performance, followed by Trem2*R47H (avg. AUC = 0.917), APOE4Trem2 (avg. AUC = 0.912), and APOE4 KI (avg. AUC = 0.910). The Elastic net model had the highest AUC (avg. AUC = 0.927), then random forest (avg. AUC = 0.918), random forest with Boruta filter (avg AUC = 0.917), and logistic regression (avg. AUC = 0.908), respectively. There were different features deemed important and selected by the feature selection methods from the final models **(Fig. 2),** but we conducted further analysis using the best performing model based on the features selected by elastic net.

**Fig. 2.**
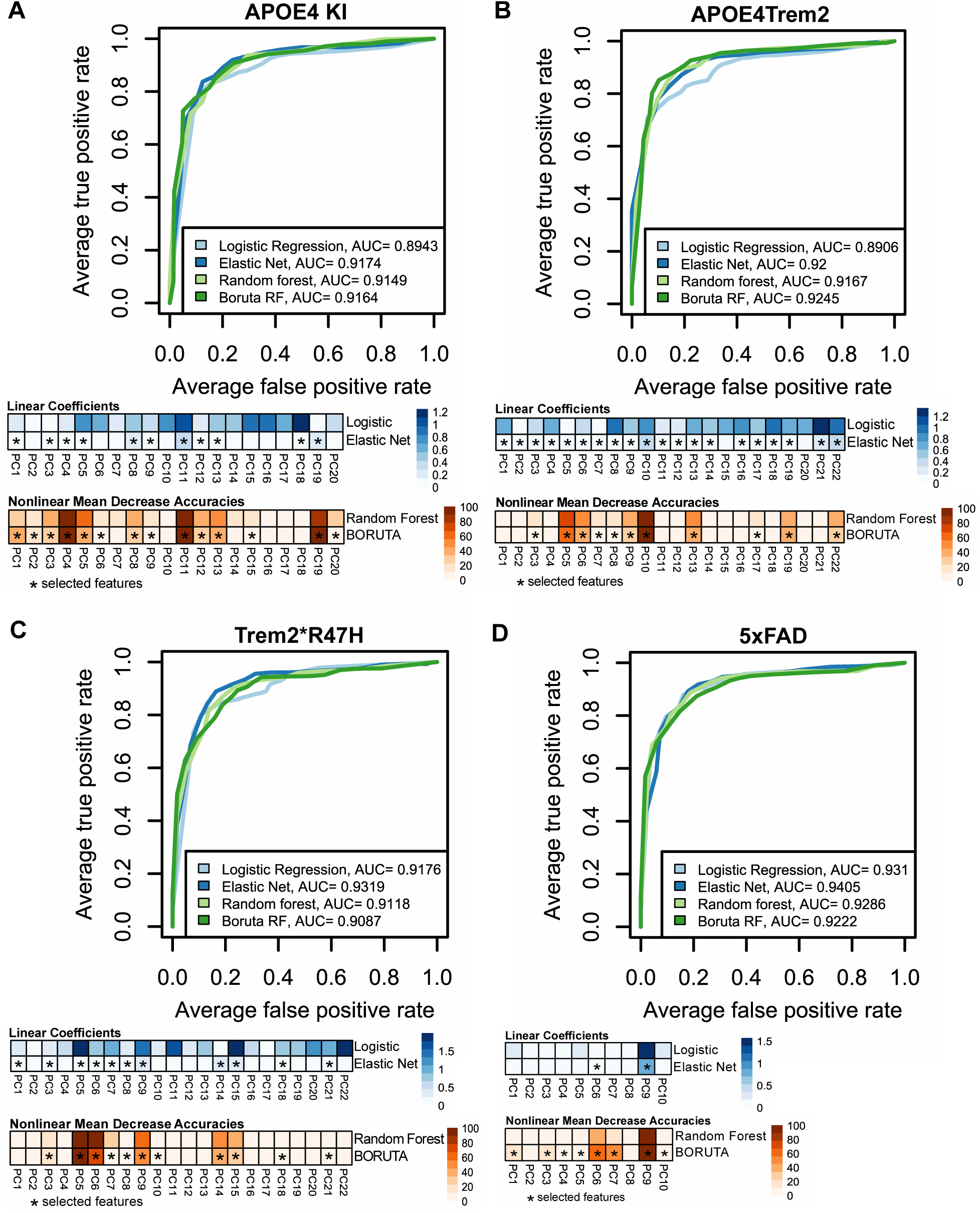
Performance for each mouse type and computational model. Averaged ROC curve across cross-validation folds predicting human AD status and the magnitudes of the coefficients (logistic regression and elastic net) and the nonlinear mean decrease accuracies (Random Forest and RF with Boruta filter) for TransCompR models using **(A)** APOE4KI. **(B)** APOE4Trem2. **(C)** Trem2*R47H. **(D)** 5xFAD. The (*) denotes features that were selected by elastic net or Boruta.

### Mouse-Derived Principal Components Explain Variance in Human AD

A strength of our cross-species modeling framework is the ability to quantify how signal in murine models predicts and explains human disease variability. We computed the mouse PC-to-human cross-species variance explained for each model’s PCs that were chosen by elastic net **(Fig. 1).** The APOE4 KI mouse PC4 had the highest variance explained in humans (50.45%) followed by APOE4Trem2 PC5 (49.47% variance explained), Trem2*R47H PC6 (13.88% variance explained), and 5xFAD PC6 (8.97% variance explained) **(Fig. 3).** In addition, each of the mouse PCs were able to clearly distinguish human AD subjects from the control group. To determine the main mouse signatures driving the differentiation between the disease condition and the healthy control in humans, we conducted fast gene set enrichment analysis (fGSEA) to identify biological pathways (Hallmark) enriched for each selected PC’s gene loadings. We identified multiple pathways enriched in human AD and shared among all the translatable mouse PCs **(Fig. 3)**. The complement pathway was enriched in APOE4 KI, APOE 4Trem2, Trem2*R47H, and 5xFAD mice. The leading-edge genes in the complement pathway shared among the mice models included *FN1, CTSS, C1QC, and C1QA*. In addition, other inflammatory pathways, including coagulation *(C1QA, FN1, A2M, SERPING1, LAMP2, PROS1, FURIN, PECAM1)*, interferon gamma response *(B2M, LATS2, SERPING1, CD74, VAMP8, STAT3),* and allograft rejection *(CTSS, B2M, CD74)* pathways were enriched in two mouse models. Additional pathways of interest included the fatty acid metabolism pathway *(HADHB, ACAA2, ADIPOR2, ALDH9A1, DECR1, UROS, GSTZ1, HSDL2, CPT1A, ENO2, HSP90AA1)* in Trem2*R47H and the estrogen response early pathway (*SLC2A1, TGM2, HSPB8, RAB31, PODXL, AFF1*) in 5xFAD AD subjects.

**Fig. 3.**
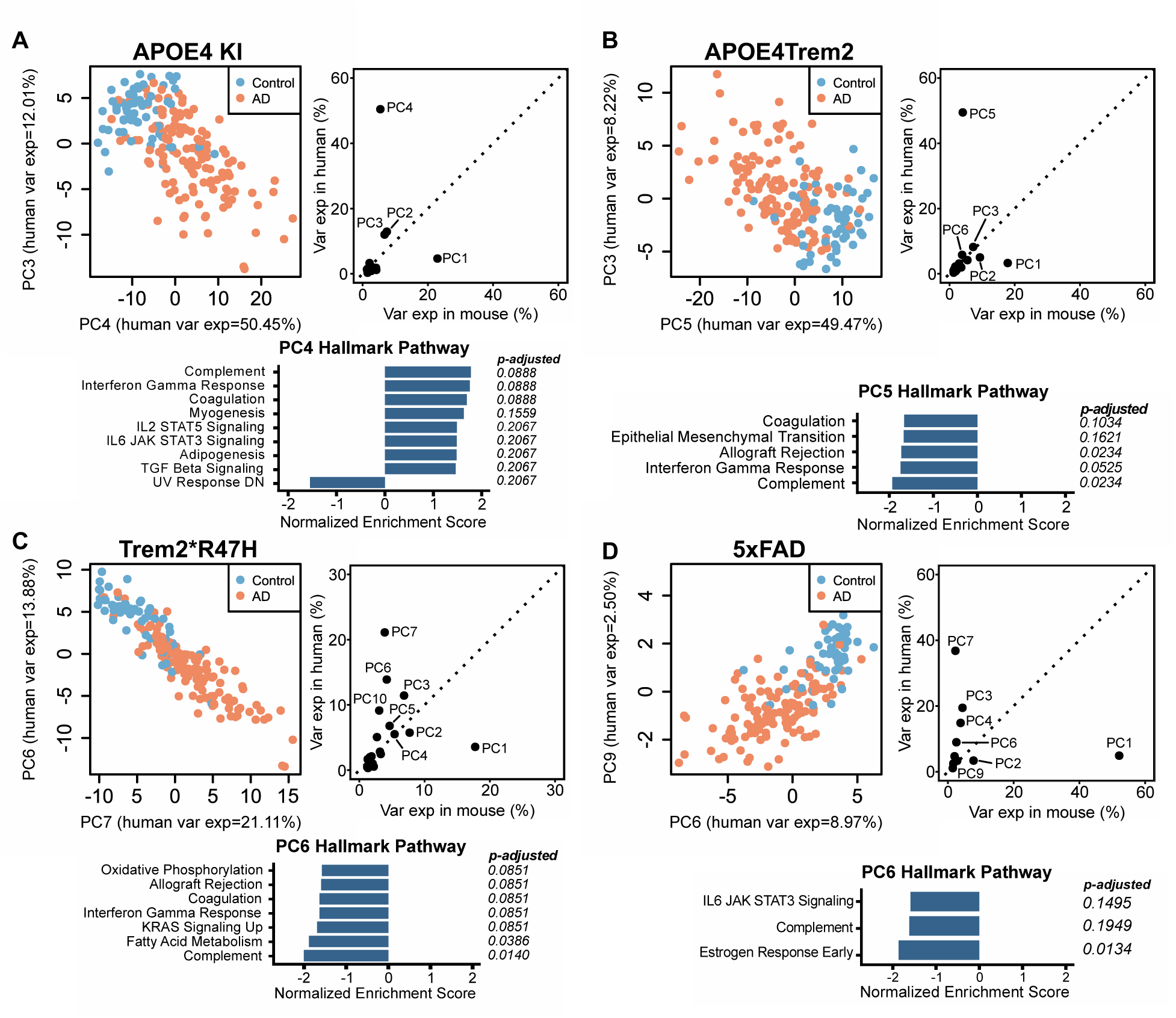
(A) Score plots for human AD subjects projected unto mouse model PCs and gene set enrichment analysis of the translatable mouse PCs. Score plot showing separation between AD and control human subjects from APOE4 KI mice PCs with high variance explained in humans. GSEA bar plots of enriched Hallmark pathways for PC4 with their normalized enrichment scores and respective adjusted p-values by Benjamini-Hochberg correction (q < 0.25). **(B)** APOE4Trem2 and GSEA results for PC5. **(C)** Trem2*R47H and GSEA results for PC6. **(D)** 5xFAD and GSEA results for PC6.

### In silico drug screening analysis identifies drugs for insomnia treatment as potential therapeutic candidates for AD

Having identified cross-species mouse-human predictive PCs, we performed an unbiased computational drug screening analysis to derive therapeutic hypotheses based on gene signatures enriched for drug response on translatable PCs. We previously developed a drug screening analysis for TransComp-R to identify PC-enriched therapeutic signatures using the LINCS L1000 database. We retrieved the characteristic direction coefficients of drugs that had defined targets based on the metadata (n=2558) from the L1000 Consensus Signature Coefficient Tables (n=33609). We computed the Spearman correlation between each drug’s differentially regulated gene coefficients and the loadings of each translatable PCs identified by TransComp-R and applied Benjamini Hochberg (B-H) FDR correction (q < 0.25) to the results. We also denoted drugs that were FDA-approved from the *Approved Drug Products with Therapeutic Equivalence Evaluations* (Orange Book, March 2025) as high value drug repurposing candidates (*21*).

The top two drugs negatively correlated with the AD group from PC4 for APOE4 KI were sotalol (*ADRB1, ADRB2, KCNH2*), an adrenergic receptor antagonist, and ramelteon (*MTNR1A, MTNR1B*), a melatonin receptor agonist. Drugs positively correlated with the disease signatures of APOE4 KI were mestinon (*ACHE*) a cholinesterase inhibitor, and sapropterin (*PAH, TH*) a phenylalanine 4-hydroxylase stimulant **(Fig. 4A).** For APOE4Trem2 translatable PCs, drugs with positive association with the control group were sotalol and liothyronine (*THRB, THRA*) a thyroid hormone stimulant. Conversely, negatively correlated drugs with the healthy subjects were voriconazole (*CYP2C19, CYP3A4, CYP2C9, PTGS1, CYP3A5*), a cytochrome P450 inhibitor, and pyrazinamide (*FASN*), a fatty acid synthase inhibitor **(Fig. 4B).** Finally, drugs positively correlated with the control group for Trem2*R47H translatable PCs included suvorexant (*HCRTR2, HCRTR1*), a dual orexin receptor antagonist, and tolterodine (*CHRM4, CHRM3, CHRM1, CHRM2*), an acetylcholine receptor antagonist **(Fig. 4C).** For 5xFAD, drugs positively correlated with the healthy group consisted of cladribine (*ADA*), an adenosine deaminase inhibitor and ribonucleoside reductase inhibitor, and nadolol (*ADRB1, ADRB2*), an adrenergic receptor antagonist **(Fig. 4D).**

**Fig. 4.**
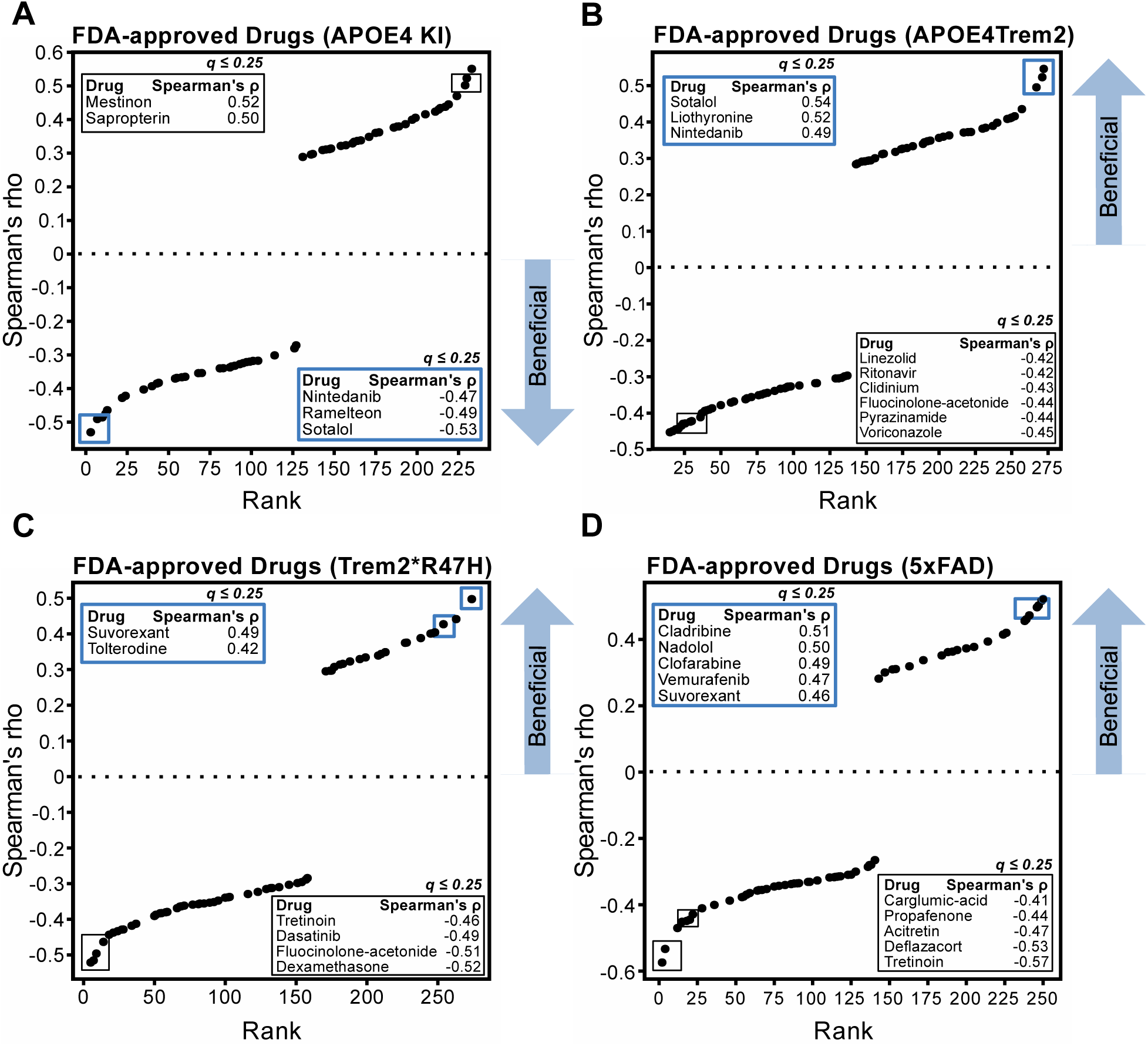
Spearman’s correlation between mouse loadings and FDA-approved drug signatures. **(A)** Ranked Spearman’s rho-values of the correlations between APOE4KI mouse loadings and the characteristic direction coefficients of each FDA-approved drug from the LINCS database. The blue boxes represent drugs with opposite signatures to the AD group. In addition, the drugs were filtered based on Benjamini-Hochberg adjusted p-values < 0.25. **(B)** APOE4Trem2. **(C)** Trem2*R47H. **(D)** 5xFAD.

Potentially therapeutic signatures were plotted to highlight genes significantly regulated by the drug and enriched in the AD or control group for each PC loading of each mouse model. We used the Protein Analysis Through Evolutionary Relationships (PANTHER) knowledgebase to determine protein classes of the gene signatures (*22*). We plotted select drug-gene to mouse-PC-gene relationships based on the biological significance of the drugs identified by statistical enrichment analysis in Figure 4 and multiple drugs were identified as potentially therapeutic for AD **(Fig. 4)**.

Ramelteon, an insomnia drug, was identified as a potentially therapeutic drug on APOE4 KI PC4. Genes downregulated in PC4 APOE4 KI AD group, such as membrane traffic proteins (*EHD3, DNM1*), and scaffold and adaptor proteins (*MICAL2, CYFIP2*), were also upregulated by ramelteon **(Fig. 5A).** Also, liothyronine, a drug used for hypothyroidism, was identified as potentially therapeutic drug on APOE4Trem2 PC5. Upregulated genes in PC5 APOE4Trem2 AD subjects, including scaffold and adaptor proteins (*SORBS1, CD81, C1QA, KCTD12, C1QB, TYROBP*), were downregulated by liothyronine **(Fig. 5B).** In addition, suvorexant, another medication for insomnia, was identified as a therapeutic candidate drug on Trem2*R47H PC6. Genes upregulated in PC6 Trem2*R47H AD group, including scaffold and adaptor proteins (*CD63, TYROBP*), were downregulated by suvorexant while downregulated genes, such as membrane trafficking regulatory proteins (*SNCA, SYT1*), were upregulated by the drug **(Fig. 5C).** Furthermore, nadolol, a drug used for high blood pressure, was identified as potential therapeutic drug on 5xFAD PC6. Genes upregulated in PC6 5xFAD AD subjects included scaffold and adaptor proteins (*SORBS1, MICAL2, KCTD12, TYROBP*) while actin or actin-binding cytoskeletal proteins (*MLC1, MYH9*) were downregulated by nadolol **(Fig. 5D).** Together, these results demonstrate significant enrichment of drugs regulating the sleep-wake cycle, thyroid function, and blood pressure as potentially translatable therapeutics for AD derived from our cross-species TransComp-R model analysis.

**Fig 5.**
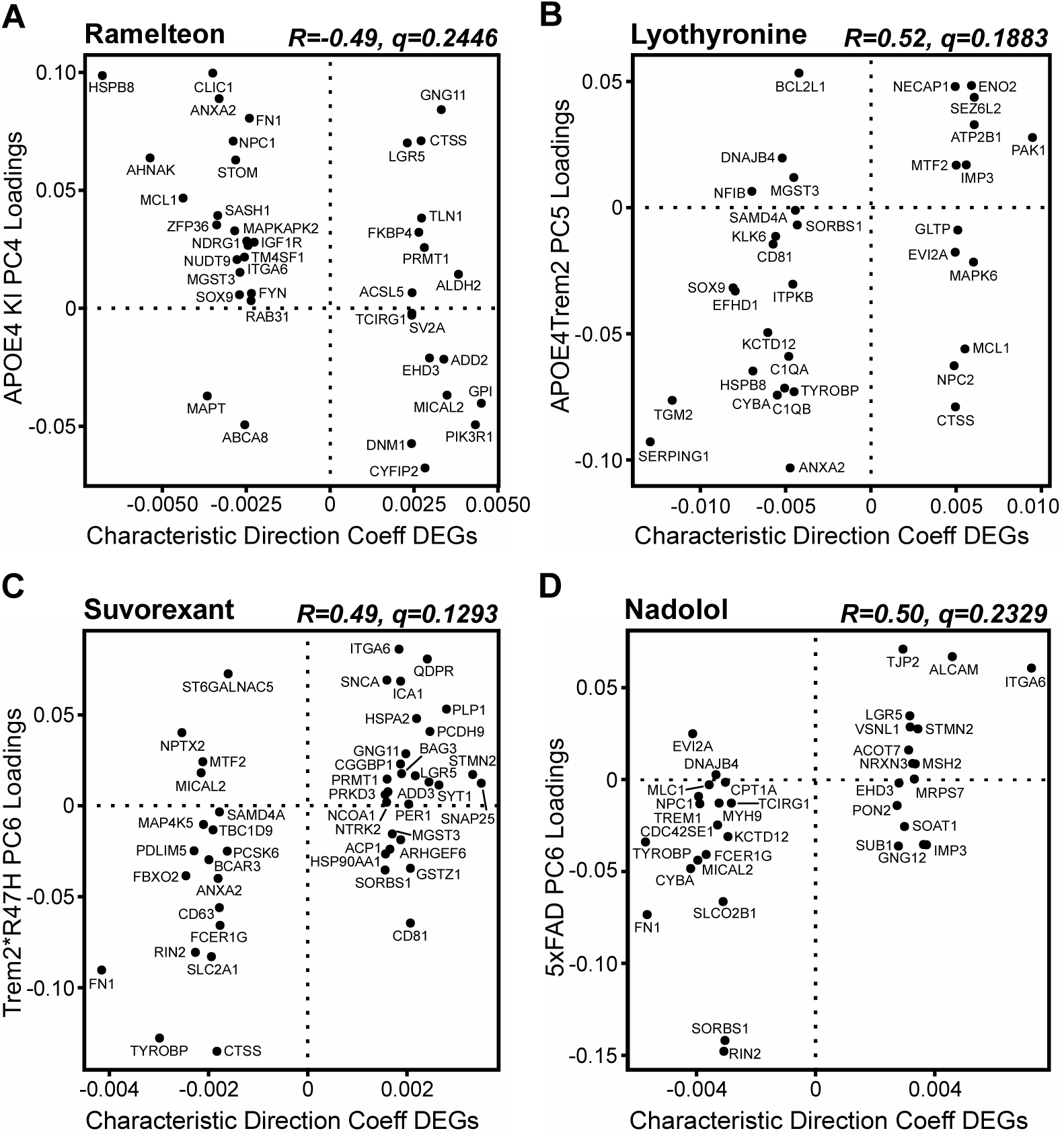
Correlation signatures for each mouse model and candidate drugs. **(A)** APOE4KI PC4 loadings and characteristic direction coefficients of DEGs from ramelteon, having a negative correlation with AD signatures. Positive characteristic direction coefficients represent genes upregulated by the drug, while negative coefficients represent downregulation. **(B)** Positive association with the control group between APOE4Trem2 PC5 loadings and liothyronine signatures. **(C)** Positive association with control between Trem2*R47H PC6 loadings and suvorexant signatures. **(D)** Positive association with the control subjects between 5xFAD PC6 loadings and nadolol signatures.

### Suvorexant Identified by TransComp-R Reduces AD Biomarkers and Modulates the CSF Proteome in an Independent Human Study

Having shown through cross-species analysis of retrospective data that FDA-approved drugs may be translatable therapeutics for AD, we tested if one of those drugs, suvorexant, could be translated in an independent human study. In a previously reported trial (*23*), participants aged 45 to 65 years with no neurological or sleep disorders were administered 20 mg of suvorexant or placebo and 6 milliliters of CSF was collected every 2-hours for 36 hours starting at 20:00. Suvorexant reduced AD biomarkers Aβ and the ratio of phosphorylated tau-181 to unphosphorylated tau-181 (pT181/T181) over hours (*23*) with the maximum effect at approximately hour 16 (12:00). Then, we performed proteomics on the same trial participants at hours 0 (baseline), 16, and 24. Proteins identified in the TransComp-R screen showed significant changes in suvorexant treated individuals. Breast cancer anti-estrogen resistance protein 3 (BCAR3), [F-actin]-monooxygenase (MICAL2), serine threonine protein kinase D3 (PRKD3), and CD63 antigen protein abundance levels had significant percent changes from baseline to t=16 compared to placebo **(Fig. 6A).** We also identified BCAR3, MICAL2, and fibronectin (FN1) protein abundance level percent changes from baseline to t=24 to be significantly different between the treatment groups **(Fig. 6B).** Suvorexant significantly decreased pT181/T181 at t=16 compared to the placebo group **(Fig. 6C).** Finally, we performed linear regression to find a potential association between AD biomarker levels and protein levels based on treatment **(Table 2).** The protein gamma-adducin (ADD3) that we identified from our cross-species model showed a significant positive linear relationship with pT181/T181 levels at t=16 in participants treated with suvorexant 20 mg **(Fig. 6D).**

**Fig. 6.**
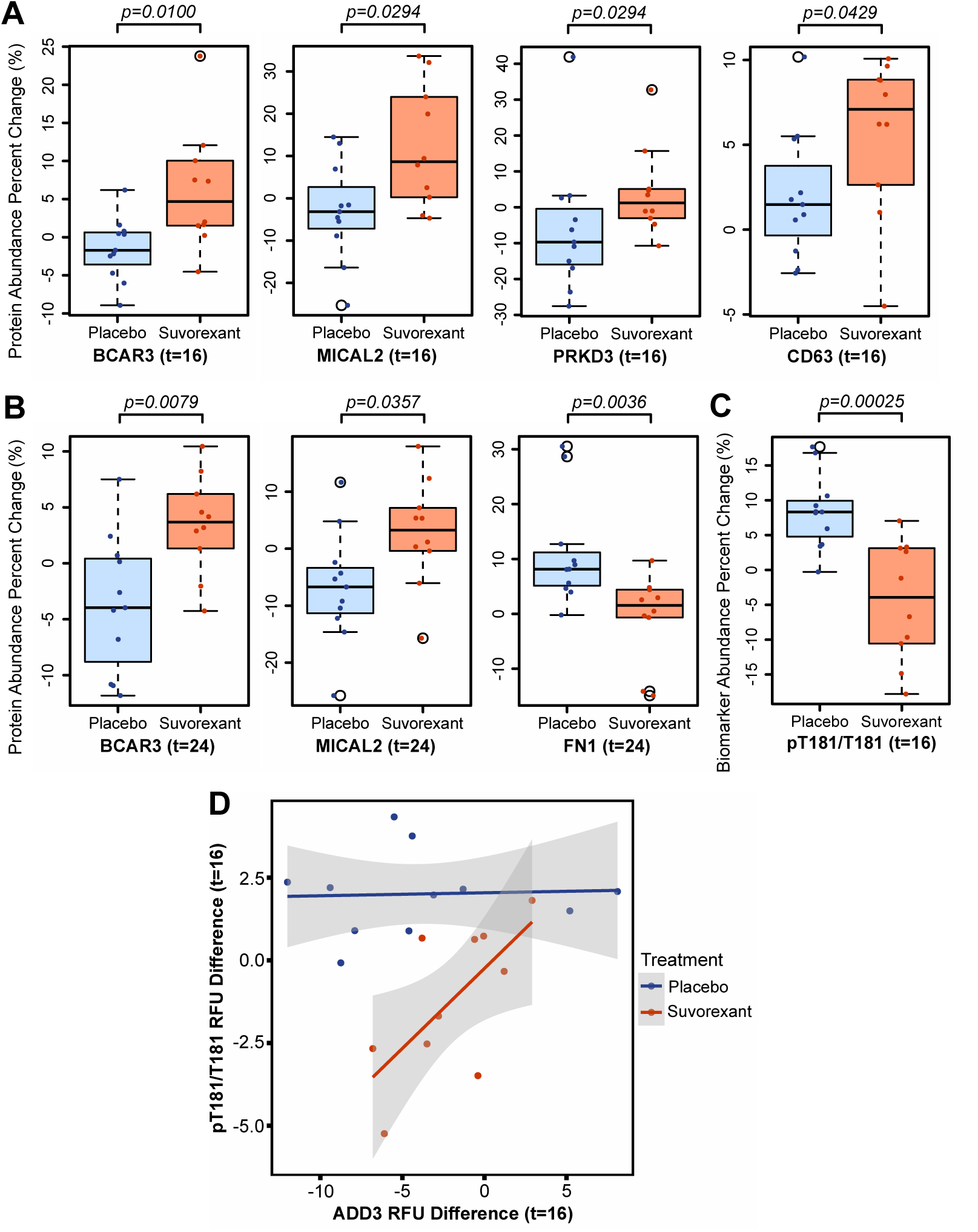
Percent changes of protein and biomarker abundance levels in CSF after treatment with suvorexant or placebo. **(A)** Percent change at t=16 from t=0 of abundance levels of proteins in the CSF that were significantly different between suvorexant and control groups, determined by Wilcoxon rank sum test (p < 0.05). **(B)** Protein abundance level percent changes in the CSF for t=24 from t=0. **(C)** Significant percent change at t=16 from t=0 protein abundance levels for pT181/T181 between suvorexant and placebo groups. **(D)** Association between the difference of ADD3 protein abundance levels at t=16 from t=0 and pT181/T181 levels based on treatment group.

**Table 2.**
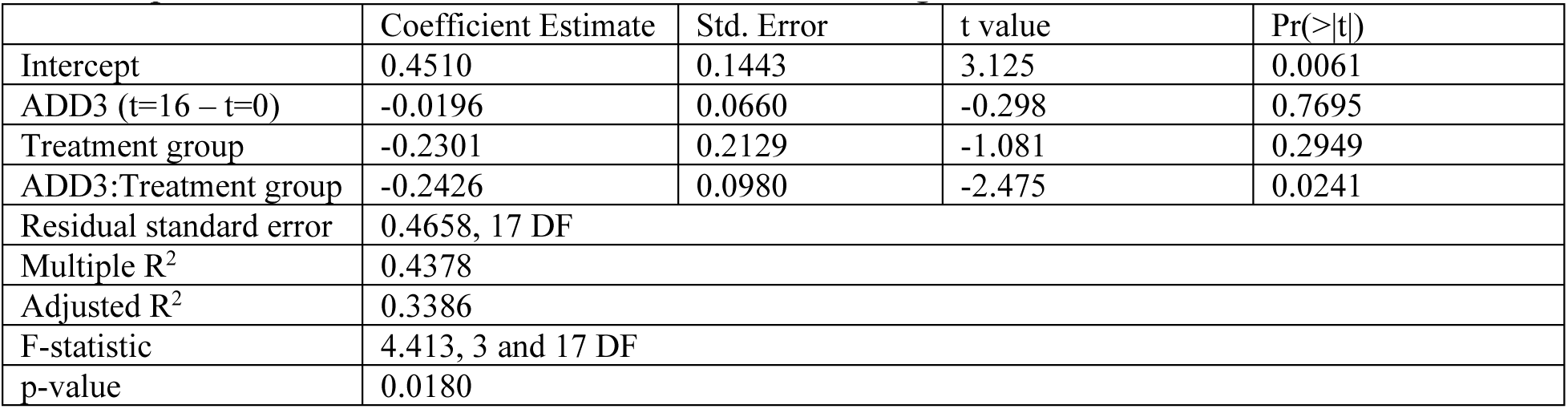
Linear regression results to determine association between ADD3 and treatment group with AD pT181/T181 biomarker at t=16 while accounting for t=0.

## DISCUSSION

In April 2025, the Food and Drug Administration (FDA) outlined a plan in conjunction with the National Institutes of Health (NIH) to reduce the overall reliance of researchers on animal models in preclinical research (*24*). Some of the alternative approaches suggested by the agencies include experimental systems (“New Advanced Methodologies” NAMS) based on human cell types, as well as computational methods for translating data from experimental studies to humans. These approaches are designed to address the lack of replicability and translatability of therapies from animal models to humans and herald a new paradigm for drug discovery and preclinical modeling of human disease.

Despite the promises of NAMS for recapitulating human biology and important regulatory steps being taken by FDA and NIH in this direction, animal models offer many unique advantages for recapitulating complex physiological processes in neuroscience and immunology. Since it is unlikely that humanized experimental systems will completely replace animal models, there is a critical need for strategies to enhance the translatability and utility of existing animal models for complex human diseases such as Alzheimer’s disease. It is in this space that computational modeling could provide flexibility and a new avenue of improving therapeutic development by integrating molecular characteristics from mouse models that can be predictive of human disease biology and therapeutic response (*18*, *25–28*). Furthermore, successfully bridging animal and human biology with computational methods in an experimental system-agnostic manner will advance these computational translation techniques for applications the inevitable discrepancies that will exist between next-generation NAMS and human disease contexts.

In this study, we used cross-species computational modeling of data from mouse models and human AD samples to identify predictive relationships between AD animal models and human disease biology. Of the four modeling methods we compared, all achieved high performance, but elastic net had the highest average AUC scores for predicting human phenotypes across multiple different mouse models (*29*). Elastic net promotes sparsity while accounting for correlated features (*30*). We coupled PC selection by elastic net with computed cross-species variance explained for each PC to identify PCs with strong predictive power and data explainability. We noted that different features were selected by RF with Boruta filtering from elastic net, potentially because different mouse model biology is emphasized by the linear and non-linear implementations of TransComp-R model (*31*). Furthermore, differences in “important” features of RF Boruta from those of linear modeling, could be due to possible inflation of mean decrease accuracy (MDA) scores due to correlated PCs (*32*).

A common feature on mouse PCs predictive of human AD was enrichment of complement signaling (*33–35*). APOE interacts with C1q to prevent the classical complement cascade from inducing synapse loss (*36*, *37*). In addition, TREM2 preserves synapses by getting involved with C1q to halt the classical complement cascade. However, decreased synapse count has been reported due to heterozygous TREM2 R47H microglia (*38*, *39*). We also found the interferon gamma response to be upregulated in AD on the translatable APOE4 KI, APOE4Trem2, and Trem2*R47H mouse PCs. Damaging effects of T cells releasing interferon-γ has been shown in mice expressing tau accumulation (*40*).

Our *in silico* drug screening identified FDA-approved drugs associated with treating sleep disorders to be potential AD therapeutics. For example, ramelteon, an insomnia drug, inversely correlated with human AD gene signatures was found to prevent activation of astrocytes and their release of pro-inflammatory cytokines in other studies (*41*, *42*). Another drug nadolol, used for high blood pressure, has low permeability across the BBB, but research has shown potential promise in its use by administering nadolol and clenbuterol in cognitively impaired subjects to promote blood flow in the brain (*43–45*). Suvorexant, a dual orexin receptor antagonist (DORA) approved by the FDA for insomnia treatment, was the most promising candidate identified in our analysis. DORAs have been shown to reduce AD hallmark biomarkers in both preclinical mouse models and humans (*23*, *46*, *47*). We identified significant protein level changes in cross-species model biomarkers BCAR3, MICAL2, PRKD3, CD63, and FN1 the treatment groups for t=16 and t=24 hours.

A novel finding of our study is that ADD3 levels may modulate the efficacy of suvorexant for reducing CSF pT1181/T181 levels. Previous research has revealed that upregulation of ADD3 1-357 fragment is associated with an increase in phosphorylated tau (*48*). Future work should further investigate the effects of suvorexant on ADD3 levels that could potentially impact phosphorylated tau levels for AD treatment and for the potential of ADD3-targetted adjuvants for enhancing suvorexant efficacy in reducing pT181/T181 levels.

The study had strengths and limitations. Our computational model was able to integrate molecular information across species and test the predictivity of the features from different mouse models in human AD through novel implementation of feature selection and nonlinear models. We determined elastic net to have increased performance through robust nested cross validation. In addition, we found multiple inflammatory pathways previously reported to be dysregulated in AD in humans that are translatable from the mouse models with potential interaction with late-onset AD genetic risk factors. Furthermore, we identified a novel signature associated with suvorexant’s role in decreasing levels of an AD biomarker in the CSF through our unbiased drug screening framework that targeted the translatable mouse features from TransComp-R. Limitations of our study, however, consisted of reduced number of genes for downstream analysis due to the mouse NanoString platform. Nevertheless, the NanoString platform provided us with a more focused selection of features from mice interpretable in humans with AD. In addition, our models did not account for demographic information of human subjects to evaluate the predictive performance of the mouse PCs. For instance, age and sex differences between control (mean age 68.6 years, 22.4% female) and AD (mean age 80.34 years, 52.3% female) human subjects available for analysis may confound our findings. Future work should expand our findings by exploring for potential age and sex differences as well as implications of other demographic factors affecting AD in humans.

Together, these results demonstrate the power of computational NAMS and TransComp-R in particular for therapeutic discovery that enhance the utility of animal models in AD research and has the potential to more rapidly evaluate drugs to repurpose to treat new diseases, expanding treatment options at a lower cost to patients.

## MATERIALS AND METHODS

### Mouse and Human Data Selection and Processing

We accessed the Gene Expression Omnibus database through *GEOquery (2.72.0), Biobase (2.64.0)* in *RStudio (R 4.4.1)* to obtain NanoString gene expression data of APOE4 KI, APOE4Trem2, Trem2*R47H, 5xFAD, and C57BL/6J mouse models of the brain hemisphere (GSE141509) (*19*, *49*, *50*). For each mouse model, we filtered by age for 6- and 12-months old to depict later stages of the disease progression. 5xFAD mouse models at these ages have demonstrated to have similar inflammatory expressions to humans (*19*). In addition, APOE4 KI and APOE4Trem2 mouse models have significant changes in gene expression from control around 6- to 9-months old (*19*). We converted the genes from the mice into human orthologous genes through *orthogene (1.10.0)* and we z-score normalized their gene expressions (*51*). For humans, we used postmortem prefrontal cortex bulk microarray gene expression data of late-onset AD (LOAD) and non-dementia subjects with ages greater than 60 years (GSE44772) (*20*). The human dataset also included demographic information, such as sex, postmortem interval (PMI), pH, and RNA integrity number (RIN) **(Table 2).** We aggregated duplicated human genes based on median scores and standardized expression values. We used genes that were shared between the mouse and human datasets for our downstream analysis.

**Table 2a.**
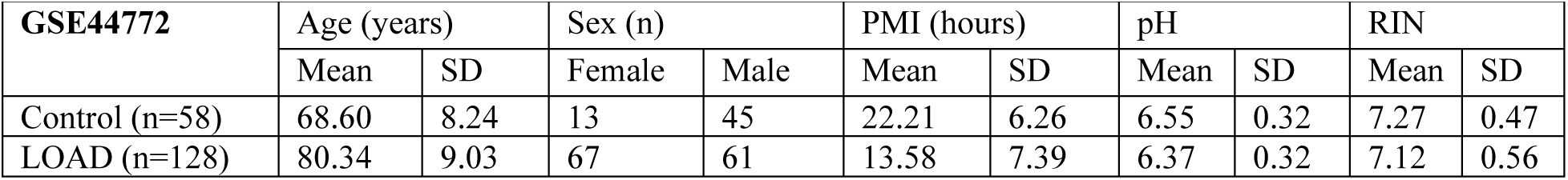
Reproduced demographic information of the human dataset from GSE44772 (*20*).

### Mouse to Human Translation through Translatable Components Regression

We performed principal component analysis on each of the mouse datasets using *prcomp* function and *factoextra (1.0.7)* for visualization for Translatable Components Regression (TransComp-R) (*52*). We selected the principal components that encompassed ∼80% of the mice variance for each mouse model and conducted matrix multiplication with human data to project human subjects into the mouse PC space, multiplying the human samples x genes matrix by the murine genes x PC’s loadings matrix. We computed the mouse variances explained in humans based on the following equation:

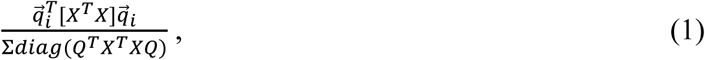

where each individual mouse PC loading i, was denoted by 𝑞 _𝑖_, Q for the matrix of all mouse PC loadings, and X for the human gene expression data.

### Linear and Nonlinear Cross-species Translation Modeling and Feature Selection

We applied logistic regression (glm) for each of the mouse models with human data integration to predict AD states using a 10-fold cross validation through *caret (6.0.94)* and obtained their averaged area under the curve values (AUC) (*53*). Similarly, we used the *caret* and *nestedcv (0.7.12)* to perform a nested 10 inner- and 10 outer-fold cross validation (CV) for hyperparameter tuning in elastic net *glmnet (4.1-8)* for feature selection (*54*, *55*). For nonlinear modeling, we ran a random forest *randomForest (4.7-1.1)* through *caret* and *nestedcv* with 10-inner and outer folds (*56*). In addition, we implemented Boruta *(8.0.0)* feature selection filtering for our random forest model through the nested CV (*31*). For the models that we conducted *nestedcv*, we retrieved the average area under the curve values of the outer folds using *ROCR (1.0.11)* (*57*). For APOE4 KI, the final model tuned parameters for elastic net were alpha=0.55 and lambda=0.0554, random forest (mtry=2), RF with Boruta (mtry=2). For APOE4Trem2, elastic net parameters were alpha=0.1 and lambda=0.0607, random forest (mtry=2), and RF with Boruta filter (mtry=2). For Trem2*R47H, parameters for elastic net were alpha=0.55, and lambda=0.0600, random forest (mtry=12), RF with Boruta (mtry=12). For 5xFAD, elastic net parameters were alpha=1, and lambda=0.0662, random forest (mtry=6), RF with Boruta filter (mtry=2).

### Gene Set Enrichment Analysis of Predictive Mouse Features

We used fast gene set enrichment analysis *fgsea (1.30.0)* with KEGG Legacy and Hallmark pathways, accessed from the Human Molecular Signatures Database, with minimal gene size of 5, maximum size of 500, and GSEA parameter of 1 (*58–60*). Statistically significant pathways were those with nominal p < 0.05 and FDR-adjusted p-adjusted value (Benjamini-Hochberg correction) q < 0.25, indicating a 25% false positive probability.

### Computational Drug Screening Analysis with LINCS L1000 Dataset

We accessed the Library of Integrated Network-Based Cellular Signatures (SigCom LINCS) for the L1000 Consensus Signature Coefficient Tables (Level 5) with gene symbols and their characteristic direction coefficients (CD) (*61–63*). We extracted the perturbagens with existing targets based on the small molecules metadata. In addition, we obtained the differentially expressed genes (DEGs) for each drug by first determining their z-scores and respective p-values from the CD coefficients and filtering genes with p-values less than 0.05 (*64*). For the CD coefficients, the signs denoted upregulation (positive) or downregulation (negative) of the genes due to the drug perturbation. We screened for drugs that could potentially mimic or reverse late-onset AD signatures by finding the Spearman correlation between each of the mouse PC loading values and the DEGs’ CD coefficients for each drug, based on overlapping genes. Then, we ranked the correlations and adjusted the p-values using Benjamini-Hochberg (q < 0.25).

### Human CSF Collection

Twenty-five cognitively unimpaired participants aged 45-65 years old were randomized to placebo or suvorexant 20 mg and underwent 36 hours of longitudinal collection of cerebrospinal fluid (CSF). As previously described, 6 ml of CSF was collected every 2 hours from 20:00 on Day 1 to 08:00 on Day 3 (*65*). Aβ isoforms (Aβ38, Aβ40, and Aβ42) and unphosphorylated and phosphorylated tau (-181, -202, -217) were measured by mass spectrometry (*65*). Participants in the suvorexant 20 mg showed decreased CSF Aβ and phosphorylated tau-181 levels compared to placebo.

### CSF Suvorexant Protein Abundance Analysis

Human CSF samples treated with suvorexant 20 mg or placebo from t=0 (20:00 on Day 1), t=16 (12:00 on Day 2), and t=24 (20:00 on Day 2) timepoints were assayed using the SOMAscan 11K kit, which is a high-scale aptamer-based proteomic platform that measures relative concentrations of ∼11,000 proteins. All CSF samples were analyzed per manufacturer’s instructions. Briefly, we inputted 50 μL of sample and prepared at a single 20% dilution across all sample wells on the SOMAscan assay plate. Samples were hybridized into Agilent Surescan MicroArray slides and then scanned at a 5-μm resolution to detect CY3 fluorescence. Gridding and image analysis were done using Agilent Feature Extraction v10.7.3.1. Raw signals were standardized according to SOMAscan (*66*).

We used the dataset with protein abundance levels from human CSF samples treated with suvorexant 20 mg (n=12) or placebo (n=13) from t=0 (20:00 on Day 1), t=16 (12:00 on Day 2), and t=24 (20:00 on Day 2) timepoints. We preprocessed the data by aggregating protein levels based on repeating subjects and proteins by taking the median. After preprocessing and excluding participants without all timepoints (t=0, 16, 24), we had (n=10) in the suvorexant group and (n=11) subjects in the placebo group. Then, we filtered the proteins based on coinciding gene symbols from our drug screening analysis. We calculated the difference for t=16 and t=24 protein abundance levels from t=0 levels and we performed Wilcoxon rank sum tests (p < 0.05) to determine significant changes of protein abundance levels of suvorexant from the placebo group. Results were visualized through boxplots of the percent change of relative protein abundance levels for t=16 and t=24 from t=0 levels.

A similar analysis was conducted for AD biomarkers, including Aβ38, Aβ40, Aβ42, T181, pT181, pT181/T181 levels, where Wilcoxon rank sum tests were performed to determine significant differences between the treatment groups, visualized through boxplots. In addition, we determined proteins associated with changes in AD biomarker levels due to suvorexant treatment through linear regression (lm). A linear regression model was built for each biomarker and each protein identified from our cross-species computational model and screening for t=16 and t=24. We modeled the AD biomarker level differences for each timepoint from t=0 as the response variable and protein difference levels from t=0, treatment group and an interaction term between the protein levels and treatment group, as independent variables. We selected models with a significant F-statistic value (p-value < 0.05) and a significant interaction between the protein and treatment group.

## FUNDING

Open Philanthropy (JHP, DKB, BPL)

Good Ventures Foundation (JHP, DKB, BPL)

Start-up funds from Weldon School of Biomedical Engineering at Purdue University (JHP, DKB) Start-up funds from Case Western Reserve University (JHP, DKB)

Washington University Personalized Medicine Initiative 2 (BPL)

## AUTHOR CONTRIBUTIONS

Conceptualization: JHP, BPL, DKB

Methodology: JHP, BPL, DKB, JY

Software: JHP

Formal analysis: JHP

Investigation: BPL, JY

Visualization: JHP

Funding acquisition: BPL, DKB

Project administration: BPL, DKB

Resources: BPL, DKB

Writing – original draft: JHP

Writing – review & editing: JHP, BPL, DKB, JY

## DATA AND MATERIALS AVAILABILITY

Mouse and human transcriptomics data are publicly available from Gene Expression Omnibus (GEO), GSE141509 and GSE44772. For the drug screening analysis, we obtained the signatures from SigCom LINCS under L1000 Consensus Signatures Coefficient Tables (Level 5) and metadata from LINCS Small Molecules Metadata (https://maayanlab.cloud/sigcom-lincs/#/Download). CSF AD biomarker and proteomic data is available to qualified investigators upon request. All code used for the analysis can be accessed at https://github.com/Brubaker-Lab/LOAD_Mouse_TransCompR.

## REFERENCES

1. K. B. Rajan, J. Weuve, L. L. Barnes, E. A. McAninch, R. S. Wilson, D. A. Evans, Population estimate of people with clinical Alzheimer’s disease and mild cognitive impairment in the United States (2020–2060). Alzheimer’s & Dementia 17, 1966–1975 (2021).

2. A. Atri, The Alzheimer’s Disease Clinical Spectrum. Medical Clinics of North America 103, 263–293 (2019).

3. B. G. Perez-Nievas, A. Serrano-Pozo, Deciphering the Astrocyte Reaction in Alzheimer’s Disease. Front. Aging Neurosci. 10, 114 (2018).

4. D. V. Hansen, J. E. Hanson, M. Sheng, Microglia in Alzheimer’s disease. Journal of Cell Biology 217, 459–472 (2018).

5. A. Serrano-Pozo, M. P. Frosch, E. Masliah, B. T. Hyman, Neuropathological Alterations in Alzheimer Disease. Cold Spring Harbor Perspectives in Medicine 1, a006189–a006189 (2011).

6. H. Hampel, J. Hardy, K. Blennow, C. Chen, G. Perry, S. H. Kim, V. L. Villemagne, P. Aisen, M. Vendruscolo, T. Iwatsubo, C. L. Masters, M. Cho, L. Lannfelt, J. L. Cummings, A. Vergallo, The Amyloid-β Pathway in Alzheimer’s Disease. Mol Psychiatry 26, 5481–5503 (2021).

7. M. F. Mendez, Early-Onset Alzheimer Disease. Neurologic Clinics 35, 263–281 (2017).

8. R. Cacace, K. Sleegers, C. Van Broeckhoven, Molecular genetics of early-onset Alzheimer’s disease revisited. Alzheimer’s & Dementia 12, 733–748 (2016).

9. D. M. Holtzman, J. Herz, G. Bu, Apolipoprotein E and Apolipoprotein E Receptors: Normal Biology and Roles in Alzheimer Disease. Cold Spring Harbor Perspectives in Medicine 2, a006312–a006312 (2012).

10. M. V. F. Silva, C. D. M. G. Loures, L. C. V. Alves, L. C. De Souza, K. B. G. Borges, M. D. G. Carvalho, Alzheimer’s disease: risk factors and potentially protective measures. J Biomed Sci 26, 33 (2019).

11. T. Jonsson, H. Stefansson, S. Steinberg, I. Jonsdottir, P. V. Jonsson, J. Snaedal, S. Bjornsson, J. Huttenlocher, A. I. Levey, J. J. Lah, D. Rujescu, H. Hampel, I. Giegling, O. A. Andreassen, K. Engedal, I. Ulstein, S. Djurovic, C. Ibrahim-Verbaas, A. Hofman, M. A. Ikram, C. M. Van Duijn, U. Thorsteinsdottir, A. Kong, K. Stefansson, Variant of *TREM2* Associated with the Risk of Alzheimer’s Disease. N Engl J Med 368, 107–116 (2013).

12. R. Guerreiro, A. Wojtas, J. Bras, M. Carrasquillo, E. Rogaeva, E. Majounie, C. Cruchaga, C. Sassi, J. S. K. Kauwe, S. Younkin, L. Hazrati, J. Collinge, J. Pocock, T. Lashley, J. Williams, J.-C. Lambert, P. Amouyel, A. Goate, R. Rademakers, K. Morgan, J. Powell, P. St. George-Hyslop, A. Singleton, J. Hardy, *TREM2* Variants in Alzheimer’s Disease. N Engl J Med 368, 117–127 (2013).

13. F. Leng, P. Edison, Neuroinflammation and microglial activation in Alzheimer disease: where do we go from here? Nat Rev Neurol 17, 157–172 (2021).

14. M. Z. Zhong, T. Peng, M. L. Duarte, M. Wang, D. Cai, Updates on mouse models of Alzheimer’s disease. Mol Neurodegeneration 19, 23 (2024).

15. J. L. Jankowsky, H. Zheng, Practical considerations for choosing a mouse model of Alzheimer’s disease. Mol Neurodegeneration 12, 89 (2017).

16. D. Sun, W. Gao, H. Hu, S. Zhou, Why 90% of clinical drug development fails and how to improve it? Acta Pharmaceutica Sinica B 12, 3049–3062 (2022).

17. D. K. Brubaker, D. A. Lauffenburger, Translating preclinical models to humans. Science 367, 742–743 (2020).

18. D. K. Brubaker, M. P. Kumar, E. L. Chiswick, C. Gregg, A. Starchenko, P. N. Vega, A. N. Southard-Smith, A. J. Simmons, E. A. Scoville, L. A. Coburn, K. T. Wilson, K. S. Lau, D. A. Lauffenburger, An interspecies translation model implicates integrin signaling in infliximab-resistant inflammatory bowel disease. Sci. Signal. 13, eaay3258 (2020).

19. the MODEL-AD Consortium, C. Preuss, R. Pandey, E. Piazza, A. Fine, A. Uyar, T. Perumal, D. Garceau, K. P. Kotredes, H. Williams, L. M. Mangravite, B. T. Lamb, A. L. Oblak, G. R. Howell, M. Sasner, B. A. Logsdon, G. W. Carter, A novel systems biology approach to evaluate mouse models of late-onset Alzheimer’s disease. Mol Neurodegeneration 15, 67 (2020).

20. B. Zhang, C. Gaiteri, L.-G. Bodea, Z. Wang, J. McElwee, A. A. Podtelezhnikov, C. Zhang, T. Xie, L. Tran, R. Dobrin, E. Fluder, B. Clurman, S. Melquist, M. Narayanan, C. Suver, H. Shah, M. Mahajan, T. Gillis, J. Mysore, M. E. MacDonald, J. R. Lamb, D. A. Bennett, C. Molony, D. J. Stone, V. Gudnason, A. J. Myers, E. E. Schadt, H. Neumann, J. Zhu, V. Emilsson, Integrated Systems Approach Identifies Genetic Nodes and Networks in Late-Onset Alzheimer’s Disease. Cell 153, 707–720 (2013).

21. Approved Drug Products with Therapeutic Equivalence Evaluations (U.S. Food & Drug Administration, ed. 45th; https://www.fda.gov/drugs/drug-approvals-and-databases/approved-drug-products-therapeutic-equivalence-evaluations-orange-book).

22. P. D. Thomas, M. J. Campbell, A. Kejariwal, H. Mi, B. Karlak, R. Daverman, K. Diemer, A. Muruganujan, A. Narechania, PANTHER: A Library of Protein Families and Subfamilies Indexed by Function. Genome Res. 13, 2129–2141 (2003).

23. B. P. Lucey, H. Liu, C. D. Toedebusch, D. Freund, T. Redrick, S. L. Chahin, K. G. Mawuenyega, J. G. Bollinger, V. Ovod, N. R. Barthélemy, R. J. Bateman, Suvorexant Acutely Decreases Tau Phosphorylation and Aβ in the Human CNS. Annals of Neurology 94, 27–40 (2023).

24. U.S. Food & Drug Administration, “FDA Announces Plan to Phase Out Animal Testing Requirement for Monoclonal Antibodies and Other Drugs” (2025); https://www.fda.gov/news-events/press-announcements/fda-announces-plan-phase-out-animal-testing-requirement-monoclonal-antibodies-and-other-drugs.

25. L. Suarez-Lopez, B. Shui, D. K. Brubaker, M. Hill, A. Bergendorf, P. S. Changelian, A. Laguna, A. Starchenko, D. A. Lauffenburger, K. M. Haigis, Cross-species transcriptomic signatures predict response to MK2 inhibition in mouse models of chronic inflammation. iScience 24, 103406 (2021).

26. B. K. Ball, E. A. Proctor, D. K. Brubaker, “Cross-Species Modeling Identifies Gene Signatures in Type 2 Diabetes Mouse Models Predictive of Inflammatory and Estrogen Signaling Pathways Associated with Alzheimer’s Disease Outcomes in Humans” in Biocomputing 2025 (WORLD SCIENTIFIC, Kohala Coast, Hawaii, USA, 2024; https://www.worldscientific.com/doi/10.1142/9789819807024_0031), pp. 426–440.

27. B. K. Ball, J. H. Park, A. M. Bergendorf, E. A. Proctor, D. K. Brubaker, Translational disease modeling of peripheral blood identifies type 2 diabetes biomarkers predictive of Alzheimer’s disease. npj Syst Biol Appl 11, 58 (2025).

28. K. M. Pullen, R. Finethy, S.-H. B. Ko, C. J. Reames, C. M. Sassetti, D. A. Lauffenburger, Cross-species transcriptomics translation reveals a role for the unfolded protein response in Mycobacterium tuberculosis infection. npj Syst Biol Appl 11, 19 (2025).

29. H. Zou, T. Hastie, Regularization and Variable Selection Via the Elastic Net. Journal of the Royal Statistical Society Series B: Statistical Methodology 67, 301–320 (2005).

30. T. J. Hastie, R. Tibshirani, J. H. Friedman, The Elements of Statistical Learning: Data Mining, Inference, and Prediction (Springer, New York, 2nd ed., 2009) Springer series in statistics.

31. M. B. Kursa, W. R. Rudnicki, Boruta: Wrapper Algorithm for All Relevant Feature Selection, (2009); 10.32614/CRAN.package.Boruta.

32. M. Rotari, M. Kulahci, Variable selection wrapper in presence of correlated input variables for random forest models. Quality & Reliability Eng 40, 297–312 (2024).

33. S. Hong, V. F. Beja-Glasser, B. M. Nfonoyim, A. Frouin, S. Li, S. Ramakrishnan, K. M. Merry, Q. Shi, A. Rosenthal, B. A. Barres, C. A. Lemere, D. J. Selkoe, B. Stevens, Complement and microglia mediate early synapse loss in Alzheimer mouse models. Science 352, 712–716 (2016).

34. B. Dejanovic, T. Wu, M.-C. Tsai, D. Graykowski, V. D. Gandham, C. M. Rose, C. E. Bakalarski, H. Ngu, Y. Wang, S. Pandey, M. G. Rezzonico, B. A. Friedman, R. Edmonds, A. De Mazière, R. Rakosi-Schmidt, T. Singh, J. Klumperman, O. Foreman, M. C. Chang, L. Xie, M. Sheng, J. E. Hanson, Complement C1q-dependent excitatory and inhibitory synapse elimination by astrocytes and microglia in Alzheimer’s disease mouse models. Nat Aging 2, 837–850 (2022).

35. B. Dejanovic, M. A. Huntley, A. De Mazière, W. J. Meilandt, T. Wu, K. Srinivasan, Z. Jiang, V. Gandham, B. A. Friedman, H. Ngu, O. Foreman, R. A. D. Carano, B. Chih, J. Klumperman, C. Bakalarski, J. E. Hanson, M. Sheng, Changes in the Synaptic Proteome in Tauopathy and Rescue of Tau-Induced Synapse Loss by C1q Antibodies. Neuron 100, 1322–1336.e7 (2018).

36. C. Yin, S. Ackermann, Z. Ma, S. K. Mohanta, C. Zhang, Y. Li, S. Nietzsche, M. Westermann, L. Peng, D. Hu, S. V. Bontha, P. Srikakulapu, M. Beer, R. T. A. Megens, S. Steffens, M. Hildner, L. D. Halder, H.-H. Eckstein, J. Pelisek, J. Herms, S. Roeber, T. Arzberger, A. Borodovsky, L. Habenicht, C. J. Binder, C. Weber, P. F. Zipfel, C. Skerka, A. J. R. Habenicht, ApoE attenuates unresolvable inflammation by complex formation with activated C1q. Nat Med 25, 496–506 (2019).

37. B. Stevens, N. J. Allen, L. E. Vazquez, G. R. Howell, K. S. Christopherson, N. Nouri, K. D. Micheva, A. K. Mehalow, A. D. Huberman, B. Stafford, A. Sher, A. M. Litke, J. D. Lambris, S. J. Smith, S. W. M. John, B. A. Barres, The Classical Complement Cascade Mediates CNS Synapse Elimination. Cell 131, 1164–1178 (2007).

38. L. Zhong, X. Sheng, W. Wang, Y. Li, R. Zhuo, K. Wang, L. Zhang, D.-D. Hu, Y. Hong, L. Chen, H. Rao, T. Li, M. Chen, Z. Lin, Y. Zhang, X. Wang, X.-X. Yan, X. Chen, G. Bu, X.-F. Chen, TREM2 receptor protects against complement-mediated synaptic loss by binding to complement C1q during neurodegeneration. Immunity 56, 1794–1808.e8 (2023).

39. J. Penney, W. T. Ralvenius, A. Loon, O. Cerit, V. Dileep, B. Milo, P. Pao, H. Woolf, L. Tsai, IPSC -derived microglia carrying the TREM2 R47H /+ mutation are proinflammatory and promote synapse loss. Glia 72, 452–469 (2024).

40. X. Chen, M. Firulyova, M. Manis, J. Herz, I. Smirnov, E. Aladyeva, C. Wang, X. Bao, M. B. Finn, H. Hu, I. Shchukina, M. W. Kim, C. M. Yuede, J. Kipnis, M. N. Artyomov, J. D. Ulrich, D. M. Holtzman, Microglia-mediated T cell infiltration drives neurodegeneration in tauopathy. Nature 615, 668–677 (2023).

41. A. Kuriyama, M. Honda, Y. Hayashino, Ramelteon for the treatment of insomnia in adults: a systematic review and meta-analysis. Sleep Medicine 15, 385–392 (2014).

42. S. Yang, J. Wang, D. Wang, L. Guo, D. Yu, Melatonin Receptor Agonist Ramelteon Suppresses LPS-Induced Neuroinflammation in Astrocytes. ACS Chem. Neurosci. 12, 1498–1505 (2021).

43. R. C. Heel, R. N. Brogden, G. E. Pakes, T. M. Speight, G. S. Avery, Nadolol: A Review of its Pharmacological Properties and Therapeutic Efficacy in Hypertension and Angina Pectoris. Drugs 20, 1–23 (1980).

44. S. Kalsoom, A. Zamir, A. U. Rehman, W. Ashraf, I. Imran, H. Saeed, A. Majeed, F. Alqahtani, M. F. Rasool, Clinical pharmacokinetics of nadolol: A systematic review. Clinical Pharmacy Therapeu 47, 1506–1516 (2022).

45. T. Lodeweyckx, J. De Hoon, K. Van Laere, E. Bautista, G. Rizzo, C. Bishop, E. Rabiner, R. S. Martin, A. Ford, G. Vargas, Effects on cerebral blood flow after single doses of the β_2_ agonist, clenbuterol, in healthy volunteers and patients with mild cognitive impairment or Parkinson’s disease. Br J Clin Pharmacol 90, 2638–2651 (2024).

46. J.-E. Kang, M. M. Lim, R. J. Bateman, J. J. Lee, L. P. Smyth, J. R. Cirrito, N. Fujiki, S. Nishino, D. M. Holtzman, Amyloid-β Dynamics Are Regulated by Orexin and the Sleep-Wake Cycle. Science 326, 1005–1007 (2009).

47. S. Parhizkar, X. Bao, W. Chen, N. Rensing, Y. Chen, M. Kipnis, S. Song, G. Gent, E. Tycksen, M. Manis, C. Lee, J. R. Serrano, M. E. Bosch, E. Franke, C. M. Yuede, E. C. Landsness, M. Wong, D. M. Holtzman, Lemborexant ameliorates tau-mediated sleep loss and neurodegeneration in males in a mouse model of tauopathy. Nat Neurosci 28, 1460–1472 (2025).

48. H. Yu, M. Xiong, C. Liu, D. Xia, L. Meng, Z. Zhang, The γ-Adducin 1–357 fragment promotes tau pathology. Front. Aging Neurosci. 15, 1241750 (2023).

49. Sean Davis <Sdavis2@Mail. Nih.Gov>, GEOquery, Bioconductor (2017); 10.18129/B9.BIOC.GEOQUERY.

50. V. C. R. Gentleman, Biobase, Bioconductor (2017); 10.18129/B9.BIOC.BIOBASE.

51. Brian Schilder, orthogene, Bioconductor; 10.18129/B9.BIOC.ORTHOGENE.

52. A. Kassambara, F. Mundt, factoextra: Extract and Visualize the Results of Multivariate Data Analyses, (2016); 10.32614/CRAN.package.factoextra.

53. M. Kuhn, caret: Classification and Regression Training, (2007); 10.32614/CRAN.package.caret.

54. M. J. Lewis, A. Spiliopoulou, K. Goldmann, C. Pitzalis, P. McKeigue, M. R. Barnes, nestedcv: an R package for fast implementation of nested cross-validation with embedded feature selection designed for transcriptomics and high-dimensional data. Bioinformatics Advances 3, vbad048 (2023).

55. J. Friedman, T. Hastie, R. Tibshirani, B. Narasimhan, K. Tay, N. Simon, J. Yang, glmnet: Lasso and Elastic-Net Regularized Generalized Linear Models, (2008); 10.32614/CRAN.package.glmnet.

56. L. Breiman, A. Cutler, A. Liaw, M. Wiener, randomForest: Breiman and Cutlers Random Forests for Classification and Regression, (2002); 10.32614/CRAN.package.randomForest.

57. T. Sing, O. Sander, N. Beerenwinkel, T. Lengauer, ROCR: Visualizing the Performance of Scoring Classifiers, (2005); 10.32614/CRAN.package.ROCR.

58. V. K. Mootha, C. M. Lindgren, K.-F. Eriksson, A. Subramanian, S. Sihag, J. Lehar, P. Puigserver, E. Carlsson, M. Ridderstråle, E. Laurila, N. Houstis, M. J. Daly, N. Patterson, J. P. Mesirov, T. R. Golub, P. Tamayo, B. Spiegelman, E. S. Lander, J. N. Hirschhorn, D. Altshuler, L. C. Groop, PGC-1α-responsive genes involved in oxidative phosphorylation are coordinately downregulated in human diabetes. Nat Genet 34, 267–273 (2003).

59. C. Alexey Sergushichev [Aut, fgsea, Bioconductor (2017); 10.18129/B9.BIOC.FGSEA.

60. A. Subramanian, P. Tamayo, V. K. Mootha, S. Mukherjee, B. L. Ebert, M. A. Gillette, A. Paulovich, S. L. Pomeroy, T. R. Golub, E. S. Lander, J. P. Mesirov, Gene set enrichment analysis: A knowledge-based approach for interpreting genome-wide expression profiles. Proc. Natl. Acad. Sci. U.S.A. 102, 15545–15550 (2005).

61. A. Subramanian, R. Narayan, S. M. Corsello, D. D. Peck, T. E. Natoli, X. Lu, J. Gould, J. F. Davis, A. A. Tubelli, J. K. Asiedu, D. L. Lahr, J. E. Hirschman, Z. Liu, M. Donahue, B. Julian, M. Khan, D. Wadden, I. C. Smith, D. Lam, A. Liberzon, C. Toder, M. Bagul, M. Orzechowski, O. M. Enache, F. Piccioni, S. A. Johnson, N. J. Lyons, A. H. Berger, A. F. Shamji, A. N. Brooks, A. Vrcic, C. Flynn, J. Rosains, D. Y. Takeda, R. Hu, D. Davison, J. Lamb, K. Ardlie, L. Hogstrom, P. Greenside, N. S. Gray, P. A. Clemons, S. Silver, X. Wu, W.-N. Zhao, W. Read-Button, X. Wu, S. J. Haggarty, L. V. Ronco, J. S. Boehm, S. L. Schreiber, J. G. Doench, J. A. Bittker, D. E. Root, B. Wong, T. R. Golub, A Next Generation Connectivity Map: L1000 Platform and the First 1,000,000 Profiles. Cell 171, 1437–1452.e17 (2017).

62. J. E. Evangelista, D. J. B. Clarke, Z. Xie, A. Lachmann, M. Jeon, K. Chen, K. M. Jagodnik, S. L. Jenkins, M. V. Kuleshov, M. L. Wojciechowicz, S. C. Schürer, M. Medvedovic, A. Ma’ayan, SigCom LINCS: data and metadata search engine for a million gene expression signatures. Nucleic Acids Research 50, W697–W709 (2022).

63. N. R. Clark, K. S. Hu, A. S. Feldmann, Y. Kou, E. Y. Chen, Q. Duan, A. Ma’ayan, The characteristic direction: a geometrical approach to identify differentially expressed genes. BMC Bioinformatics 15, 79 (2014).

64. Z. Xie, E. Kropiwnicki, M. L. Wojciechowicz, K. M. Jagodnik, I. Shu, A. Bailey, D. J. B. Clarke, M. Jeon, J. E. Evangelista, M. V. Kuleshov, A. Lachmann, A. A. Parigi, J. M. Sanchez, S. L. Jenkins, A. Ma’ayan, Getting Started with LINCS Datasets and Tools. Current Protocols 2, e487 (2022).

65. B. P. Lucey, H. Liu, C. D. Toedebusch, D. Freund, T. Redrick, S. L. Chahin, K. G. Mawuenyega, J. G. Bollinger, V. Ovod, N. R. Barthélemy, R. J. Bateman, Suvorexant Acutely Decreases Tau Phosphorylation and Aβ in the Human CNS. Annals of Neurology 94, 27–40 (2023).

66. J. Candia, G. Fantoni, F. Delgado-Peraza, N. Shehadeh, T. Tanaka, R. Moaddel, K. A. Walker, L. Ferrucci, Variability of 7K and 11K SomaScan Plasma Proteomics Assays. J. Proteome Res. 23, 5531–5539 (2024).

