## Supplemental Figure 1 for "Cross-Species Translation Enhances the Use of Mouse Models for Translatability and Drug Discovery in Late-Onset Alzheimer’s Disease"

### SUPPLEMENTARY INFORMATION

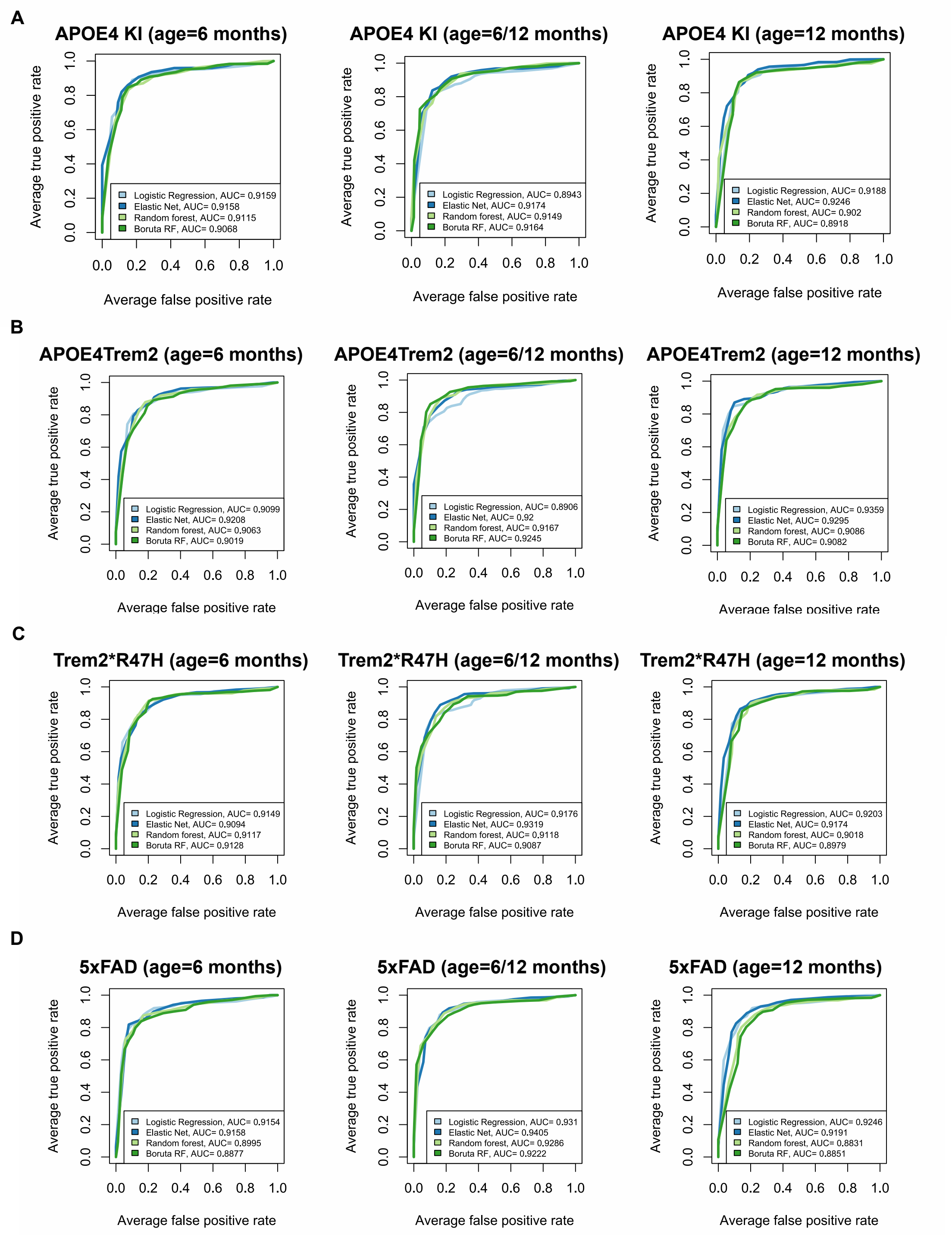


**Fig. S1. Performance for each mouse model, computational model, and age.** (A) Average AUC scores for logistic regression, elastic net, random forest, and random forest with Boruta filter performances for integrated human subjects with APOE4 KI at different mice ages, including only 6-months, 6- and 12-months, and only 12-months, from left to right. (B) AUC scores for APOE4Trem2 for different mice ages. (C) AUC scores for Trem2*R47H for different mice ages. (D) AUC scores for 5xFAD for different mice ages.
